# Time-resolved neural reinstatement and separation during memory decisions in human hippocampus

**DOI:** 10.1101/196212

**Authors:** Lynn J. Lohnas, Katherine Duncan, Werner K. Doyle, Orrin Devinsky, Lila Davachi

**Affiliations:** Department of Psychology, 6 Washington Place, 8^th^ floor, New York, NY 10003; Center for Neural Science, New York University; 6 Washington Place, 8^th^ floor, New York, NY 10003; Department of Psychology, University of Toronto; 100 St. George Street Rm 4018, Toronto, ON M5S 3G3; Department of Neurosurgery, New York University School of Medicine; 550 First Avenue, New York, NY 10016; Department of Neurology, Epilepsy Division, New York University School of Medicine; 223 East 34^th^ Street, New York, NY 10016

**Keywords:** hippocampus, electrocorticography, memory reinstatement, pattern separation

## Abstract

Mnemonic decision-making has long been hypothesized to rely on hippocampal dynamics that bias memory processing toward the formation of new memories or the retrieval of old ones. Successful memory encoding would be best optimized by pattern separation, whereby two highly similar experiences can be represented by underlying neural populations in an orthogonal manner. By contrast, successful memory retrieval is thought to be supported by a recovery of the same neural pattern laid down during encoding. Here we examined how hippocampal pattern completion and separation emerge over time during memory decisions. We measured electrocorticography activity in the human hippocampus and posterior occipitotemporal cortex (OTC) while participants performed continuous recognition of items that were new, repeated (old), or highly similar to a prior item (similar).During retrieval decisions of old items, both regions exhibited significant reinstatement of multivariate high frequency activity (HFA) associated with encoding. Further, the extent of reinstatement of encoding patterns during retrieval was correlated both with the strength (HFA power) of hippocampal encoding and with the strength of hippocampal retrieval. Evidence for encoding pattern reinstatement was also seen in OTC on trials requiring fine-grained discrimination of similar items. By contrast, hippocampal activity showed evidence for pattern separation during these trials. Together, these results underscore the critical role of the hippocampus in supporting both reinstatement of overlapping information and separation of similar events.

**Significance Statement:** One of the biggest computational challenges the memory systems faces is to disambiguate highly similar experiences while at the same time preserving and reinstating prior memories. Remarkably, hippocampal processes have been implicated in both of these functions. However, how this is accomplished is unknown. Leveraging the spatiotemporal resolution of electrocorticography, we found evidence for memory reinstatement in both the hippocampus and occipitotemporal cortex. Reinstatement was differentiated in time across these two regions with earlier reinstatement evident in occipitotemporal cortex. Interestingly, when a current experience was very similar, but not identical to a prior one, occipitotemporal cortical activity still showed reinstatement of the prior memory but hippocampal activity differentiated or disambiguated these two similar experiences.

## Introduction

Perhaps one of the most challenging functions of the episodic memory system is distinguishing between two experiences that contain highly overlapping content. Pattern separation refers to the process of representing highly similar events in a distinct way, thus allowing them to co-exist with minimal interference (2–7). Yet if two distinct experiences share overlapping features, then the second experience may also promote “pattern completion” of the first experience (2, 4, 8). These two processes – pattern completion and separation – reflect opposing if not contradictory functions, both of which have been attributed to the hippocampus. To reconcile how the hippocampus can accomplish both processes, it has been proposed that novelty may bias the hippocampal system towards pattern separation while familiarity may promote memory retrieval, or pattern completion (9–12).How and when the hippocampus can support representations of both overlapping and distinctive features of events has been an active area of research.. Nonetheless, the temporal dynamics examining how these processes emerge over the course of a single memory decision remains relatively unexplored.

Theoretical and rodent work provide evidence that the hippocampus exhibits sensitivity to differences in highly similar events while also representing the events’ strong overlap (2, 4, 13, 14). For instance, when comparing place cell firing across two similar environments, the place cell location remained the same but the firing rates differed between chambers (15). In a similar way, Knierim and colleagues (16,17) examined place cell activity between two environments with global and local cues rotated, and found that subsets of place cell locations were consistent with either the rotated local or global cue rotations, thus keeping track of the original environment location as well as the rotated cue of the new environment. However, place cell studies cannot answer whether, on a cognitive level, the rodent is successful in recognizing the similarities and differences across two environments. In addition, place cell studies typically record activity over more extended periods of time and multiple visits to the same location, and thus are unable to answer how pattern completion and separation emerge upon the second encounter with a similar or identical location. One study began to address this latter question using a context-dependent associative reward-learning task (18). They found that different subsets of hippocampal cells exhibited firing rates that distinguished between the context (i.e., the spatial environment) and an item’s identity, position or valence.Nonetheless, these hippocampal responses were recorded following an initial learning phase, and thus do not capture hippocampal dynamics during the initial phase of distinguishing between similar memories.

In humans, multivariate approaches have been used to examine mnemonic reinstatement effects for similar and identical stimuli, both with functional magnetic resonance imaging (fMRI) (19) and electrocorticography (ECoG). In the hippocampus, there is evidence for reinstatement of encoding patterns during successful later retrieval both for similar stimuli from the same category (20) and for identical stimuli (10, 21–23). However, such effects are not unique to the hippocampus. Mnemonic reinstatement in several cortical regions has been shown with ECoG (24–28), and has been noted in visual cortex (29, 30), medial temporal lobe cortex (10, 21, 31, 32), and prefrontal cortex (29) using fMRI.

At a mechanistic level, cortical reinstatement during retrieval may result from or interact with hippocampal pattern completion (33, 34). Supporting this idea, cortical reinstatement has been shown to correlate with hippocampal univariate activity both at encoding (30, 35) and retrieval (10, 21, 23, 29, 32, 36– 38). Thus, hippocampal computations may recover the memory representations formed during encoding and this may, in turn, support cortical reinstatement.Nonetheless, the fMRI response, on the time scale of seconds, is not well suited to address how reinstatement emerges over time and across regions, and to our knowledge no ECoG study has contrasted reinstatement with separation in the hippocampus.

While there is strong evidence that both cortical regions and the hippocampus contribute to memory reinstatement, multivariate fMRI studies have uniquely implicated the hippocampus in supporting pattern separation of highly similar experiences. In particular, there is evidence that hippocampal pattern separation is greater for very similar item pairs compared to unrelated item pairs (5, 6, 31, 7). In addition, studies of univariate fMRI activity provide evidence that hippocampal activation is sensitive to whether a presented item is either identical or just highly similar to a previously presented item (31, 39–46). In one seminal example (39), unlike other hippocampal subregions or MTL cortical regions, hippocampal subregion DG/CA3 did not show repetition suppression for highly similar lure items; rather, activity was not significantly different between lures and new items. These results have been interpreted as evidence that the DG/CA3 subregion of the hippocampus plays a unique and critical role in distinguishing highly overlapping memory representations (39), consistent with prior theoretical work (4, 14) and rodent work (47).

Taken together, there is accumulating evidence that hippocampal processes contribute both to memory reinstatement, or pattern completion, and to pattern separation. However, it is not understood how these distinct operations are orchestrated in time over the course of a memory decision. In particular, it is not known whether reinstatement and separation occur on similar time scales in the hippocampus, nor whether reinstatement occurs on similar time scales in cortical regions as in the hippocampus. Furthermore, it remains unclear how attentional focus on the overlapping or distinctive features of similar stimuli modulates neural reinstatement or separation (48). To address these questions, we took advantage of the spatiotemporal resolution of ECoG activity, comparing activity contributing to separation and reinstatement within individual trials and across different regions. Specifically, we recorded depth and surface cortical ECoG activity as participants performed a continuous recognition paradigm previously used to examine fine-grained mnemonic discrimination (2, 9, 49, 50). In each participant we examined dynamics of high-frequency activity (HFA; 45-115 Hz), an established correlate of firing rates of individual neurons (51–53) and of fMRI activity (54, 55). In order to directly address questions about pattern separation and completion, in addition to univariate measurements, we adopted a multivariate pattern similarity approach to measure the overlap between neural representations of presented stimuli.

We examined both univariate and multivariate HFA measures in the hippocampus, posterior occipitotemporal cortex (OTC), and dorsolateral prefrontal cortex (DLPFC). We specifically considered regions both upstream (OTC) and downstream (DLPFC) from the hippocampus to characterize the timing of the flow of mnemonic activity patterns across regions. We chose OTC because we reasoned that visual object processing regions may be sensitive to the perceptual similarity between a visually presented item and its similar prior presentation. We chose DLPFC because, much like the hippocampus, this region is necessary for maintenance of contextual aspects of episodic memory (56, 57) and studies of associative memory have consistently noted that DLPFC and hippocampal regions exhibit activity modulated by successful memory encoding (58–63) and successful memory retrieval (24, 43, 64, 65). Our experimental set-up additionally allowed us to address the extent to which reinstatement and separation are modulated by task demands. Across two blocks, participants viewed a series of objects, which could be new, repeated, or highly similar but not identical to a previously presented object. In the fine-grain task block (Fig. 1A), participants classified objects as new, old, or similar; distinguishing between the latter two categories required a fine-grained mnemonic discrimination based on the visual features of the stimuli. By contrast, in the coarse-grain task block participants were instructed to classify both old and similar items as ‘old’ (Fig. 1B).

**Fig. 1.**
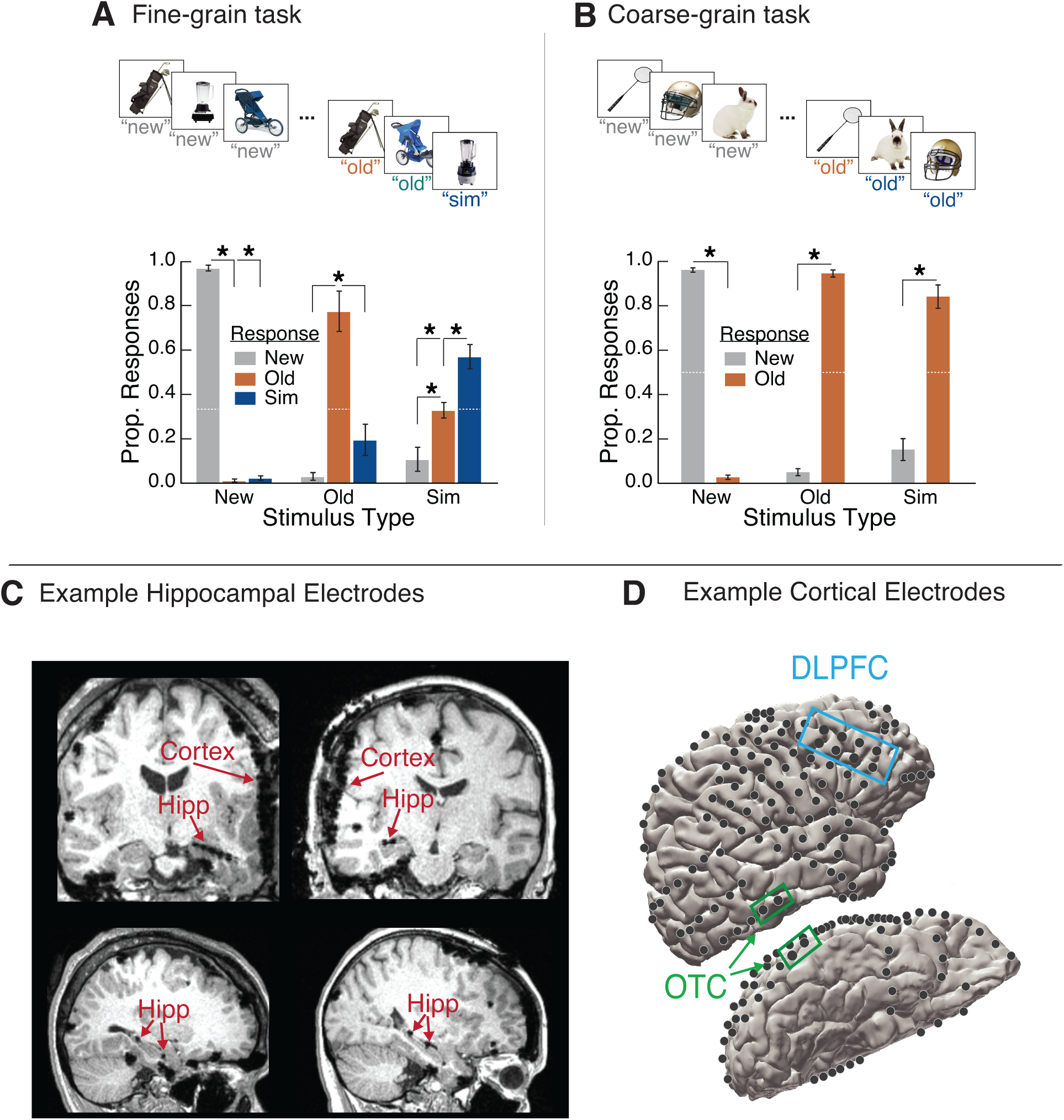
Experiment design behavioral results, and regions of interest (N=5). (A) In the fine-grain task block, participants viewed images one at a time and had to distinguish between new items (new), exact repeats (old), and items that were similar but not identical to a previous presentation (similar). Top: The four stimulus types considered in the electrophysiological data are shown here: correct new items, correct old items, correct similar items, and similar items incorrectly classified as old. Bottom: Memory performance by stimulus and response type in the fine-grain task. Error bars indicate mean ±SEM across participants. *p<.05, **p<.01.(B) In the coarse-grain task block, participants viewed new, old and similar items, but classified both similar and old items as ‘old’, thus not requiring as fine-grained discrimination between these latter two stimulus types. Top: The three stimulus types examined in the electrophysiological data are shown here: correct new items, correct old items, and correct similar items. Bottom: Memory performance by stimulus and response type in the coarse-grain task. Error bars indicate mean ±SEM across participants. *p<.05, **p<.01. (C) Hippocampal electrode placements in the hippocampus for Participant 5 (left) and Participant 1 (right). Hippocampal electrodes were visualized using each participant’s post-operative magnetic resonance imaging scan. (D) Electrode placements in posterior occipitotemporal cortex (OTC) and dorsolateral prefrontal cortex (DLPFC) for Participant 1. Cortical surface electrode placements were visualized on each participant’s rendered 3D brain (94).

## Results

### Behavior

Fig. 1A,B shows the proportion of responses as a function of stimulus type. In both tasks, memory performance was above chance (dashed white lines) for all stimulus types (fine-grain task: chance=.33; new: p<.0001, t(4)=54.5, old: p=.0057, t(4)=5.39, similar: p=.0086, t(4)=4.80; coarse-grain task: chance=.50; new: p<.0001, t(4)=53.3, old: p<.0001, t(4)=31.6, similar: p=.0019, t(4)=7.29). Further, in the coarse-grain task, participants were more likely to classify new items as new than as old (p<.0001, t(4)=54.7), and old items as old than new (p<.0001, t(4)=32.1). Similarly, in the fine-grain task, participants were more likely to classify new items correctly than to classify them as old (p<.0001, t(4)=62.5) or similar (p<.0001, t(4)=49.7).Participants were also accurate in their responses to old items in the fine-grain task,classifying them significantly more often as old items than as new (p=.0015, t(4)=7.74) or as similar (p=.0158, t(4)=4.02).

As expected, in the fine-grain task, accuracy was significantly lower for similar items in comparison to new items (p=.0016, t(4) = 7.58) and to old items (p=.0327, t(4)=3.21). However, in the coarse-grain task, the correct classification of similar items (as old) was not lower than correct classification of new items (p=.0638, t(4)=2.54) or old items (p=.0623, t(4)=2.56). This reflects the fact that in the coarse-grain task, participants did not have to discriminate between old and similar items. Indeed, if a similar item was misclassified in the fine-grain task, it was more likely to be misclassified as ‘old’ than ‘new’ (p=.0253, t(4)=3.48), further suggesting that errors for similar items in the fine-grain task primarily arose from an inability to discriminate whether the similar item was the same or slightly different from its corresponding original presentation, rather than an inability to recognize that a version of the stimulus was presented previously. Further supporting the notion that additional mnemonic discrimination was needed and deployed in the fine-grain task, response times were significantly faster in the coarse-grain task compared to the fine-grain task (p=.0036, t(4)=6.13; fine-grain mean ± SEM=1.60s±.14; coarse-grain mean ± SEM=1.37s±.11).

### Memory-related differences in univariate ECoG HFA

Our first set of analyses focused on correct trials only, across conditions and regions. HFA in both hippocampus and OTC exhibited significant time-sensitive responses in the fine-grain task (Fig. 2). To examine the temporal dynamics of memory processing in each region, we examined differences in HFA divided into 500ms time bins. First, we compared HFA for new and old presentations of a stimulus (66–69).In the hippocampus, HFA was significantly greater for new items than old items during the later 1.5-2s time window (Fig. 2A; p=.016, actual Z=20.4, null mean Z=-.283). By contrast, HFA in OTC did not exhibit significant differences between old and new items.

**Fig. 2.**
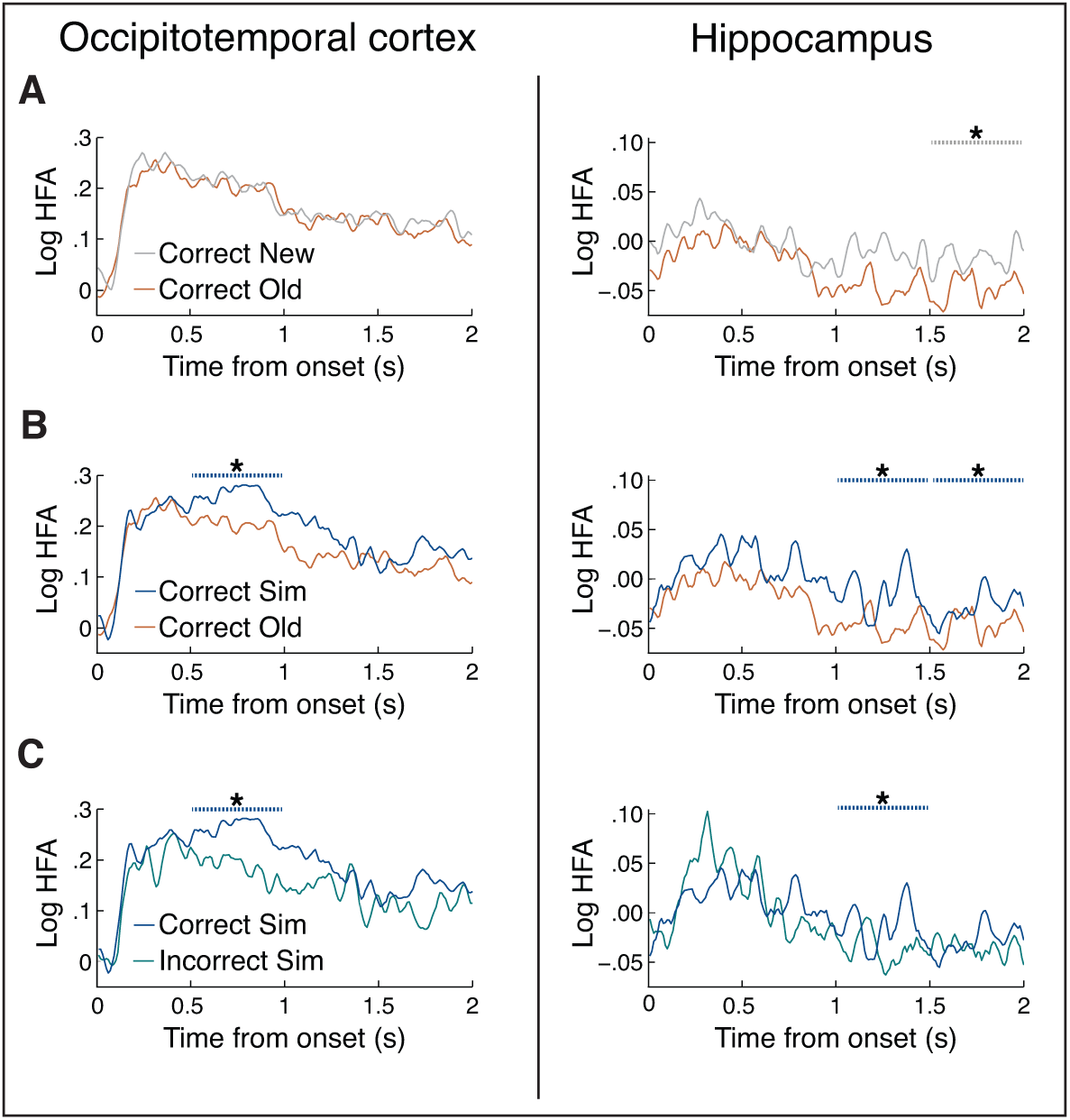
In the fine-grain task, high-frequency activity (HFA) in temporal lobe regions discriminated between stimulus and response type. Significance was assessed in the 2s following post-stimulus onset divided into four 500ms time bins, and is plotted as dashed lines above HFA. (A) HFA for correct old items and correct new items that were subsequently correctly classified as old. HFA in posterior occipitotemporal cortex (OTC) did not exhibit significant differences between old and new items, whereas hippocampal HFA was significantly reduced for old items during 1.5-2s post-stimulus onset. (B) HFA in both OTC and hippocampus was significantly greater for correct similar items than correct old items. (C) HFA in both OTC and hippocampus was significantly greater for correct similar items than similar items incorrectly classified as old. *p<.05. N=5.

We next asked whether HFA distinguishes between old items and similar items (correct sim), arguably one of the most challenging components of this task as similar items place more demands on mnemonic discrimination between the visual features of the current stimulus and one retrieved from memory. We found that HFA in both hippocampus and OTC was significantly greater for similar compared to old items (Fig. 2B). Interestingly, in OTC, this effect emerged during the early 0.5-1s time bin (p<.001, actual Z=15.8, null mean Z=.005), whereas in hippocampus this effect occurred later and lasted longer, through 1-1.5s and 1.5-2s (1-1.5s: p=.032, actual Z=16.7, null mean Z= .289; 1.5-2s: p=.048, actual Z=17.1, null mean Z=.136). Moreover, when considered with the old/correct new comparison above, hippocampal HFA was significantly greater for new items than old items during the same time window as when HFA is greater for similar items than old items, with no significant difference between similar and new items (p>.3). This pattern of activity, also seen in prior fMRI work (2, 39, 49, 70), is consistent with the notion that old items evoke stronger repetition suppression that highly similar lures. Elevated hippocampal HFA for similar relative to old items, occurring at a relatively late timepoint, may reflect attention to and/or encoding of the novel details of the similar items, thus allowing mnemonic resolution between old and similar items, consistent with the unique role of the hippocampus in separation. However, we see evidence that hippocampal processing was necessary for similar items beyond novelty detection: we find that HFA for similar and new items is not identical in all time windows: HFA for similar items was significantly greater than new items in both OTC and in hippocampus during the .5-1s time bin (p<.001, actual Z=15.1, null mean Z=-.133; hippocampus: p=.016, actual Z=19.1, null mean Z=.417).

While the above result suggests that univariate measures of HFA are sensitive to the increased mnemonic demands required to discriminate similar trials and old trials, in order to directly assess whether HFA is related to successful mnemonic discrimination, we next compared HFA during successful versus unsuccessful similar item discrimination. We hypothesized that HFA would be related to successful discrimination and thus would be significantly greater for correctly classified similar items compared to similar items incorrectly classified as ‘old’. We found that in both hippocampus and OTC, HFA was significantly greater for correct than incorrect similar items (Fig. 2C). Importantly, we found that this effect was dissociated in time: OTC exhibited a significant difference in the early 0.5-1s window (p=.024, actual Z=9.62, null mean Z=.168) and hippocampus exhibited a significant difference in a later 1-1.5s window (one-tailed p=.032, actual Z=14.5, null mean Z=.256).

To determine whether this timing was significantly different between OTC and hippocampus, we queried the difference in Z-scores between correct and incorrect similar items in each the two time bins noted above (0.5-1s minus 1-1.5s), for each brain region (OTC and hippocampus). We posited that the difference in Z-scores should be greater in OTC than hippocampus, and found that this was indeed the case (one-tailed ranksum p=.0476; OTC mean Z= 1.15; hippocampus mean Z= -.775).

In summary, analysis of mean HFA power in hippocampus and OTC suggest that activity in both regions may contribute to successful mnemonic discrimination. First, both regions exhibited HFA that distinguished between correct similar and correct old items. More directly, HFA in both regions differentiated between trials where discrimination of similar items was successful versus not, with greater HFA for successful trials than those for which similar items were classified as ‘old’.Furthermore, the fact that the HFA differences emerge in an earlier time window in OTC than in hippocampus further suggests that the hippocampus is likely receiving and incorporating information from earlier occipitotemporal cortical regions.

### Item-level reinstatement and separation

While the univariate data is suggestive of memory processes associated with pattern completion and separation, to more directly measure these processes, we next examined multivariate patterns in our data. Specifically, we calculated the similarity in HFA patterns evoked during each item’s first presentation (as a new item) and its subsequent presentation either as an old or similar item. An HFA spatiotemporal pattern (STPS) for each trial was defined as a vector of HFA power values, where a single point in the vector corresponded to HFA during a 50ms time bin at a particular electrode. Thus, for each subject, a vector of HFA values was concatenated across all electrodes within an ROI and all 50ms time bins within a 500ms time bin (see Methods for complete details). Using this vector, we then calculated the similarity between HFA patterns for an item’s first presentation and its second presentation as a similar or old item (Fig. 3A). Critically, to eliminate more general contributions to pattern similarity, we compared the similarity scores of the actual matched trial pairs (i.e., between each item’s first and second presentations) to a null distribution with shuffled trial labels (see Methods).

**Fig. 3.**
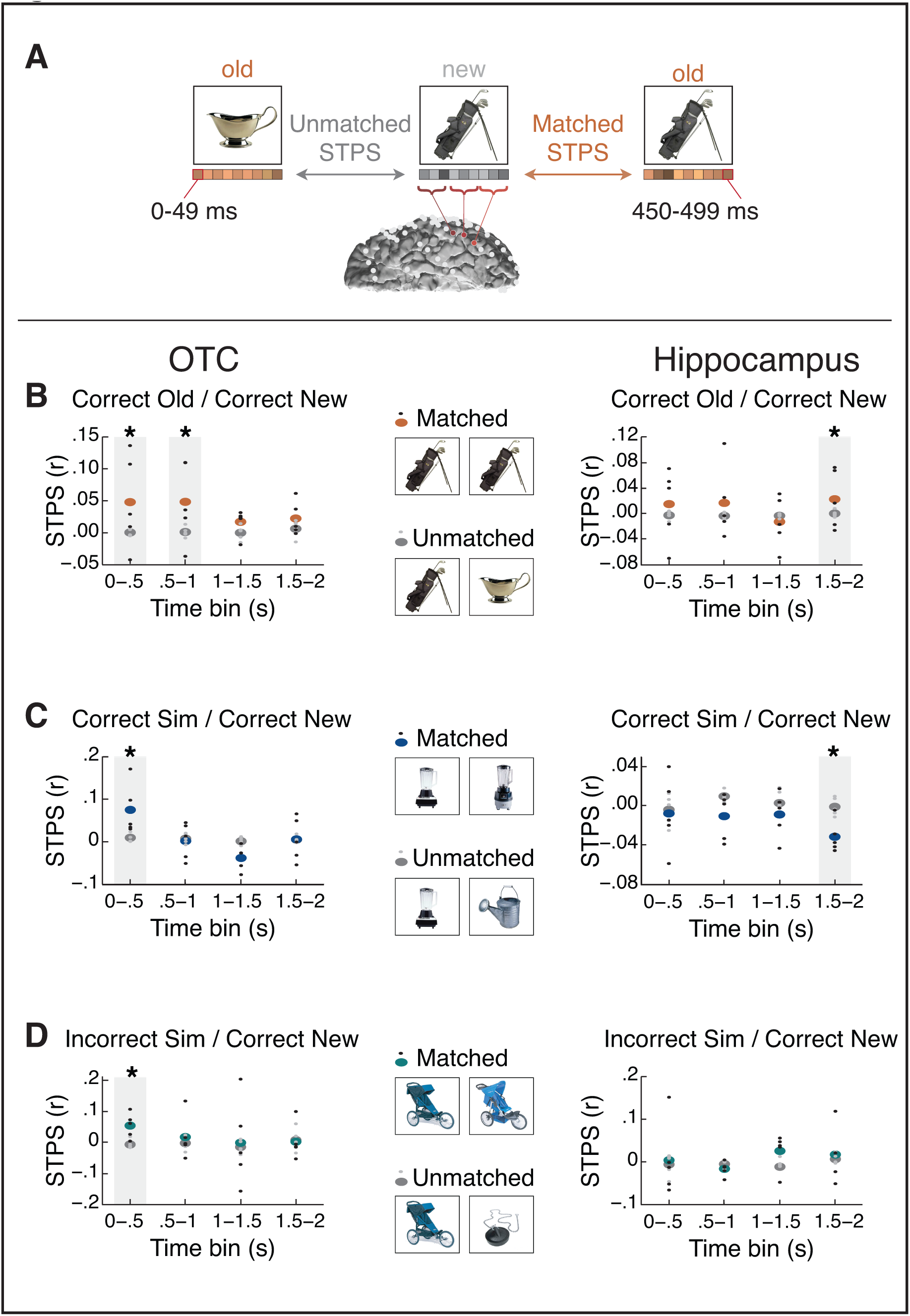
Spatiotemporal pattern similarity (STPS). (A) STPS was calculated across frequency and time and concatenated across all electrodes in each region of interest. We considered STPS between item’s matched first and second presentations, and assessed significance against a baseline of unmatched first and second presentations (see Methods for details). (B) STPS of old items in the fine-grain task. Both posterior occipitotemporal cortex (OTC) and hippocampus exhibited positive STPS, indicative of reinstatement. (C) STPS of similar items in the fine-grain task. OTC exhibited positive STPS indicative of reinstatement, whereas hippocampus exhibited significantly negative STPS, indicative of differentiation. (D) STPS of similar items classified as ‘old’ in the fine-grain task. OTC exhibited reinstatement of these items, whereas hippocampus did not exhibit significant differences. *p<.05. N=5.

We first hypothesized that correct old trials should be associated with significant reinstatement of the original encoding pattern, as measured by a greater correlation between a trial’s retrieval pattern and its encoding pattern, consistent with past findings from fMRI both in hippocampus (10, 20–23) and cortex (10, 21, 25–32). We thus computed HFA STPS between matched correct old/correct new pairs, focusing our analyses on the same time windows as when univarate HFA was significantly different for old items in these regions (Fig. 3B). We found evidence for reinstatement in both OTC and hippocampus, namely that STPS in both regions was significantly greater for matched pairs compared to the null distribution (hippocampus, 1.5-2s: one-tailed p=.03, actual mean=.0228, null mean=.0004; OTC,.5-1s: p=.015, actual mean=.0480, null mean=.0007). In OTC, this significant pattern similarity occurred in an even earlier window, as well: 0-.5s (p=.02, actual mean=.0476, null mean=.000004).

Memory reinstatement is likely to be most robust when the presented item is the same as one that was initially presented. Furthermore, strong memory reinstatement may work against mnemonic discrimination when the current item is similar, but not identical to, the original presentation. For such items, the evoked pattern should be more distinct, or pattern separated, from its original presentation to support successful mnemonic discrimination. We thus asked whether hippocampal STPS between new items and their similar presentations would be significantly *reduced*, or more distinct, than expected by chance, expecting such an effect to occur during one of the later time windows where we saw that HFA discriminated between old and similar items. Critically, during the 1.5-2s time window, the hippocampus exhibited significantly *reduced* STPS than the null distribution, providing evidence for separation in this region (Fig. 3C; one-tailed p=.045, actual mean=-.0319, null mean=-.0013). Indeed, during this time STPS was significantly greater for old-new item pairs than similar-new item pairs (Fig. 4A; p=.015, actual mean=.0548, null mean=.0017). These results provide the first evidence for pattern separation in hippocampal HFA activity patterns during mnemonic discrimination of highly similar items (1–3).

**Fig. 4.**
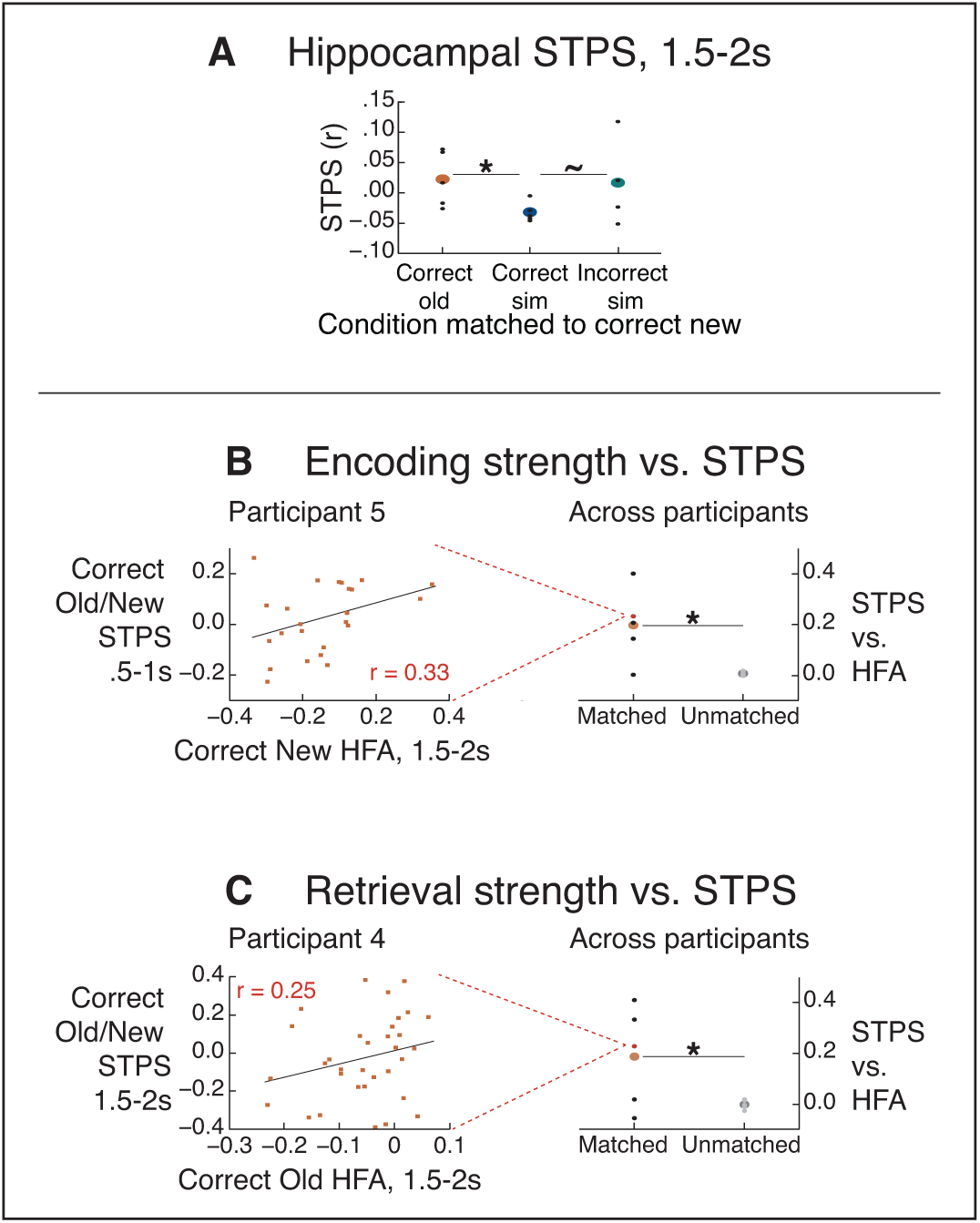
Hippocampal spatiotemporal pattern similarity (STPS) in the fine-grain task across conditions, and correlations with univariate high frequency activity (HFA). (A) In the hippocampus 1.5-2s post-stimulus onset, there was significantly more separation of similar items in comparison to correct old items, and a trend towards more separation in comparison to incorrect similar items (classified as old). (B) Left: Representative example of calculating each participant’s correlation between hippocampal encoding HFA and hippocampal STPS. Right: Across participants, hippocampal HFA during encoding of items subsequently correctly classified as old was significantly correlated with the extent of hippocampal STPS for the old items.(C) Left: Representative example of calculating each participant’s correlation between hippocampal retrieval HFA and hippocampal STPS. Right: Across participants, hippocampal HFA during retrieval of correct old items was significantly correlated with the extent of hippocampal STPS for the old items. ∼p<.1.*p<.05. N=5.

In OTC, we also queried whether there would be significant STPS effects for similar items in the same time bins that we saw reinstatement in OTC. By contrast to hippocampus, no significant separation was seen in OTC. Instead, STPS between new and similar items was significantly greater than expected by chance, consistent with reinstatement in this region, in the 0-.5s time bin (p=.020, actual mean=.0755, null mean=.0094).

Thus far, we have examined pattern similarity for correct items only. A critical step, however, is to query more directly how reinstatement and separation are related to behavioral success in mnemonic discrimination. We next examined the extent of reinstatement or separation for similar items incorrectly classified as ‘old’, limiting this analysis to trials whose first presentation was correctly classified as ‘new’. Given that in OTC we saw significant reinstatement of correct similar items 0-.5s after stimulus presentation, we anticipated that, if there were significant reinstatement of incorrect similar items, it would be during this time window as well, and indeed this is what we found: In OTC during the 0-.5s time bin, STPS was significantly greater for incorrect similar trials (incorrect sim/correct new) in comparison to a null distribution (Fig. 3D; p=.020, actual mean=.0548, null mean=-.0059). Thus, OTC exhibited reinstatement irrespective of whether the similar items were correctly or incorrectly classified, and pattern similarity between these conditions was not significant (p=.435, actual mean=.0026, null mean=-.0035). Thus this early reinstatement may be relatively automatic and stimulus evoked rather than reflect a top-down mnemonic operation.

By contrast, hippocampal STPS did not exhibit any evidence for significant reinstatement or separation between matched and unmatched pairs for incorrect similar trials. Given that the hippocampus exhibited significant pattern reinstatement for old items and pattern separation for similar items in the same 1.5-2s time window, we asked whether, in this time window, we would also see less separation for incorrect similar items in comparison to correct similar items. Thus, we compared STPS between similar items incorrectly labeled as ‘old’ (incorrect sim/correct new) and correct similar items (correct sim/correct new), and found a trend towards more separation for correct similar items (Fig. 4A; one-tailed p=.065, actual mean=.049, null mean=.0076). Thus we only see significant pattern separation in hippocampus for trials correctly identified as similar.

These pattern analyses reveal that, like univariate HFA power, both OTC and hippocampus distinguish between correct similar and old items. However, critically, OTC showed evidence only for reinstatement, whereas hippocampal patterns showed evidence for both memory reinstatement during exact repeats of old items and for pattern separation when presented with highly similar items.

### Univariate hippocampal HFA correlates with hippocampal pattern reinstatement

Theoretically, the extent of memory reinstatement during retrieval for a given item should be related to how well that item was initially encoded, i.e. its memory strength. Here we tested this assumption by asking whether hippocampal reinstatement was related to encoding strength. To this end, we quantified the encoding strength of each item as hippocampal HFA during encoding of new items (30, 35, 36) during 1.5-2s, as this was the time window with significant differences between old vs. new items. Then, for each participant, we calculated the correlation between hippocampal encoding strength and hippocampal HFA pattern similarity on a trial by trial basis for matched correct old/correct new pairs for each of the four .5s time bins from 0-2s. Indeed, we found evidence that hippocampal HFA during encoding was correlated with pattern reinstatement in the 0.5-1s time window (Fig. 4B; p=. 020, actual mean=.1986, null mean=-.0113).

We next asked if the extent of memory reinstatement is also related to the strength of unvariate hippocampal retrieval activity (10, 21, 29, 32, 36–38). As with the encoding strength, we quantified the retrieval strength as univariate hippocampal HFA for old items for the 1.5-2s time window, as hippocampal HFA exhibited significant differences for old vs. new items during this time. Although we examined the correlations for retrieval strength in all four time bins, we found a significant correlation only in the 1.5-2s time bin (Fig. 4C; p=.040, actual mean=.1818, null mean=-.0028), thus suggesting a correlation between the reinstatement of hippocampal patterns and the univariate HFA retrieval response. Thus, taken together, the results suggest that strong hippocampal encoding activity is related to early memory reinstatement, whereas later memory reinstatement seems to be more related to a strong HFA retrieval response.

### Modulation of hippocampal HFA by task demands

So far, we have shown that univariate HFA in the hippocampus and OTC exhibited significant differences between correct old, similar and new items in the fine-grain task, where task demands required discrimination between all three stimulus types. One final question we asked was whether these effects are sensitive to task demands or are more automatic in nature. To this end, we examined HFA in these same regions in the same participants while they performed a task nearly identical to the one described thus far except they did not have to distinguish between similar and old items; although participants viewed the same three stimulus types, they responded ‘old’ to both similar and old items (Fig. 1B).

Interestingly, we found that in this coarse-grain task, hippocampal univariate HFA did not exhibit any significant differences between stimulus types (Fig. 5), in contrast to what we observed for the fine-grain task. Further, the hippocampus did not exhibit significant STPS differences for matched vs. unmatched pairs. This suggests that the hippocampal contribution to memory processing and discrimination is sensitive to task demands, such that its activity discriminates between similar and old items more strongly when participants are attending to and are required to respond to these differences.

**Fig. 5.**
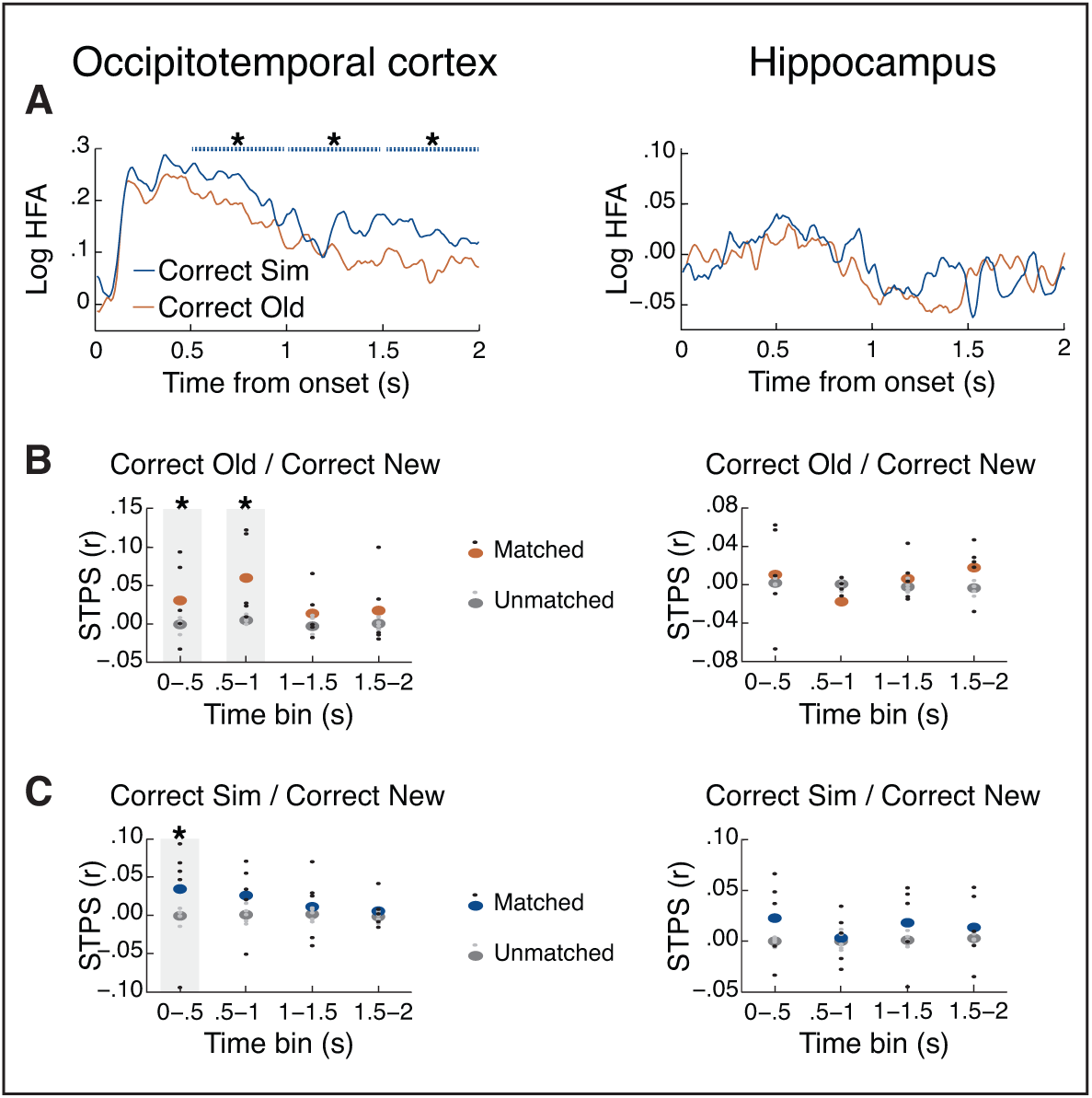
High-frequency activity (HFA) and spatiotemporal pattern similarity (STPS) of temporal lobe regions in the coarse-grain task. Significance was assessed in the 2s following post-stimulus onset divided into four 500ms time bins, and is plotted as dashed lines above HFA. (A) HFA in posterior occipitotemporal cortex (OTC) distinguished between correct old and correct similar items, despite not being relevant to task demands. Hippocampal HFA did not distinguish between stimulus types. (B) STPS of correct old items. As in the fine-grain task, OTC exhibited significant reinstatement for these items. Unlike the fine-grain task, hippocampal STPS did not exhibit any significant differences. (C) STPS of correct similar items. As in the fine-grain task, OTC exhibited significant reinstatement for these items. Unlike the fine-grain task, hippocampal STPS did not exhibit any significant differences.*p<.05. N=5.

By contrast, HFA in the OTC was not sensitive to task demands and still showed greater HFA for similar compared to old items during the 0.5-1s time window (p=.012, actual mean Z=11.4, null mean Z=-.020; note that in this version of the task, incorrect similar items to which the participant responded old are the correct similar items). Unlike the fine-grain task, this significant effect continued throughout the 2s interval (1-1.5s: p=.008, actual mean Z =10.6, null mean Z =-.021; 1.5-2s: p=.02, actual mean Z =10.4, null mean Z =-.121). In addition, we asked whether STPS in OTC would continue to exhibit significant reinstatement in the same time bins as in the fine-grain task. We found this to be the case: reinstatement for old items was significantly greater for matched vs. unmatched items during 0-.5s (one-tailed p=.04, actual mean=.0299, null mean=-.0013) and .5-1s (p<.001, actual mean=.0592, null mean=.0041), as well as for similar items during 0-.5s (p=.035, actual mean=.0352, null mean=.0001). Thus, taken together, our results show that hippocampal HFA, but not HFA in OTC, was sensitive to task demands.

### HFA in dorsolateral prefrontal cortex (DLPFC)

Thus far, our results have shown that OTC appears to be sensitive to stimulus type irrespective of task demands and hippocampal HFA is very sensitive to task demands, only distinguishing between the memory status of items in the fine-grain task. To address whether the hippocampus’ sensitivity to task demands might be a more general property of ‘higher-level’ mnemonic brain regions we examined HFA in the DLPFC. Like hippocampus, DLPFC has been implicated in successful memory encoding (58–63) and successful memory retrieval (24, 43, 64, 65).

DLPFC HFA did discriminate between stimulus types, but these dissociations were upheld irrespective of task demands (Fig. 6A). Most importantly, HFA was significantly greater for similar items than old items in the fine-grain task during the 1-1.5s and 1.5-2s time windows (1-1.5s: p=.036, actual Z=27.5, null mean Z=-.542; 1.5-2s: p=.020, actual Z=31.0, null mean Z=-.225), and significant differences in these time windows remained when considering the coarse-grain task (1-1.5s: p<.001, actual Z=36.1, null mean Z=.074; 1.5-2s: p=.030, actual Z=25.5, null mean Z=-.056). These across-task similarities in DLPFC univariate HFA contrasts with the task differences in hippocampal HFA, yet are consistent with the role of DLPFC in successful memory processing (24, 59, 71, 72). Lastly, we examined HFA pattern similarity effects in DLPFC (Fig. 6B), but there were no significant differences (fine-grain task: all p’s > .08; coarse-grain task: all p’s > .06).

**Fig. 6.**
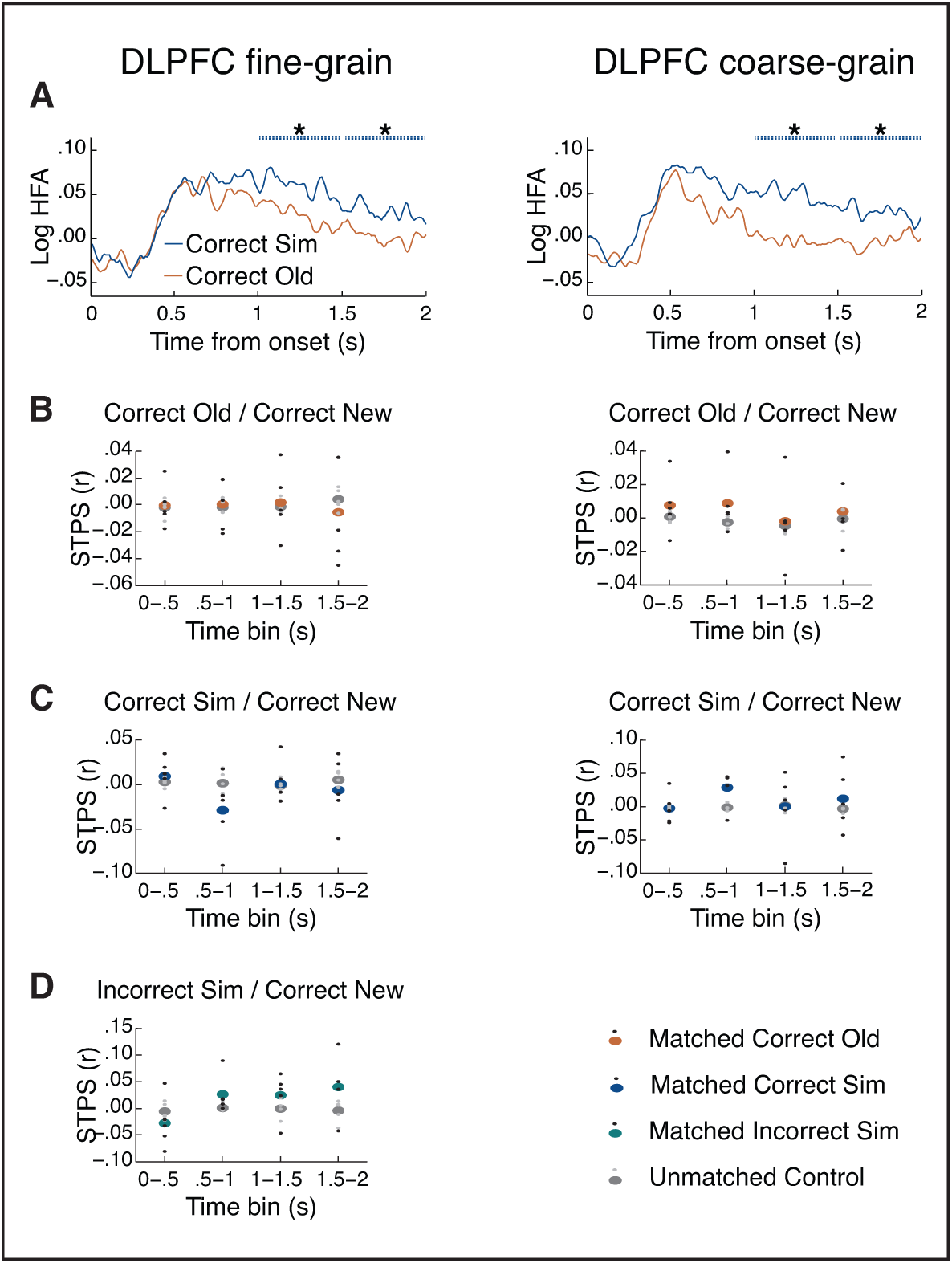
High-frequency activity (HFA) and spatiotemporal pattern similarity (STPS) in dorsolateral prefrontal cortex (DLPFC) across tasks and stimulus types. (A) DLPFC HFA was significantly greater for correct similar items than correct old items in both tasks. (B) STPS across tasks and stimulus types. *p<.05. N=5.

## Discussion

Distinguishing between two highly similar events poses a unique challenge to the episodic memory system. On the one hand, there may be an advantage to preserving access to the unique aspects of each memory. On the other hand, representing the two events’ overlapping information may also promote generalization and learning. Interestingly, hippocampal processes have been hypothesized to contribute to both of these functions – reinstating prior memory patterns and forming unique memories – but understanding how and when these processes unfold over time in the hippocampus is not known. To this end, we examined electrophysiological activity in the human hippocampus as well as in occipitotemporal cortex (OTC) and dorsolateral prefrontal cortex (DLPFC) for signatures of both memory reinstatement and separation as participants performed a paradigm requiring fine-grain mnemonic discrimination between old and similar items.

We found that univariate high frequency activity (HFA), an established measure of local neuronal firing (51–53), differentiated all memory trial types: correct new, old and similar items in the hippocampus. Most notably, hippocampal HFA was significantly enhanced for similar items compared to *both* correctly recognized old items as well as similar items incorrectly classified as old. This is consistent with prior fMRI work showing increased BOLD activation for similar trial compared to old trials (39, 41, 44, 48). Using multivariate pattern analyses of the HFA signal, we also see evidence for significant memory reinstatement when old items were presented and pattern separation when similar, but not identical, items were presented. Interestingly, hippocampal reinstatement and separation were both significant during 1.5-2s, and univariate HFA was significantly different between old and similar items during this time, as well. To our knowledge, these are some of the first reported findings examining how hippocampal separation and reinstatement emerge over time between an item’s first and second presentations, suggesting that these processes may emerge on similar time scales. Furthermore, on a trial-by-trial basis, hippocampal HFA pattern reinstatement during retrieval of old items was correlated with hippocampal HFA during both encoding and during retrieval. These results suggest that the level of memory reinstatement as measured by hippocampal encoding-retrieval similarity can be used to probe retrieval of individual items (10, 21–23), and that the strength of reinstatement correlates with univariate activity at encoding (30, 35) and at retrieval (10, 21, 23, 29, 32, 36–38).

Interestingly, the correlation between hippocampal encoding and retrieval strength and the reinstatement of multivariate memory patterns were dissociated in time: hippocampal encoding HFA correlated with early hippocampal pattern reinstatement at the .5-1s time window after stimulus presentation, whereas the hippocampal retrieval HFA correlated with hippocampal pattern reinstatement in a later time window, 1.5-2s after stimulus presentation. The brief and dissociated timing of each of these effects may help to resolve why not all fMRI studies find significant correlations at both encoding and retrieval, as these more transient effects may be difficult to measure with the slow timescale of fMRI. More crucially, this suggests that memory reinstatement may be supported by different processes at different timepoints in hippocampus – perhaps with early reinstatement being triggered by the cue strength and with later reinstatement being modulated by operations engaged during retrieval itself. Taken together, these data highlight that both encoding and retrieval processes serve to modulate reinstatement during memory decisions.

Perhaps surprisingly, all of the hippocampal effects were task dependent and were not evident in a task that presented but did not require participants to discriminate between old and similar items. In this coarse-grain task, hippocampal HFA did not show significant differences between stimulus types, either in the univariate or the pattern similarity analyses. It is unlikely that this lack of significant effects during the coarse grain task is driven by other more global factors, such as the task being easier and participants may not be as engaged because activity patterns in other regions (OTC and DLPFC, described in more detail below) did not differ as a function of the task demands. Thus, these results suggest that active attentional and goal processes need to be considered and incorporated into existing models of hippocampal pattern separation and reinstatement.

Prior ECoG work has provided evidence for hippocampal HFA activity and pattern reinstatement during memory decisions (23, 73) that engendered the recovery of contextual details. A recent study (74) examined hippocampal ECoG activity while participants performed a recognition task similar to the current fine-grain task, where they were shown images of celebrities and at test had to discriminate between highly similar pictures of viewed celebrities. Using single-unit ECoG recordings, they found *decreased* neuronal firing in CA3/DG when viewing lures (similar items) in comparison to targets (old items), and that the extent of this decrease correlated with successful memory discrimination. Given that HFA has been shown to correlate with neuronal firing (51–53), it might seem surprising that, in a similar early time window in the fine-grain task we found significantly *increased* HFA for similar than old items. However, it is possible that several methodological differences across these studies may have contributed to the disparate findings. For example, whereas we presented each stimulus only once before presenting its matching old or similar item, this prior work allowed nine study sessions with each item, and all old and similar items were tested three times (74). Furthermore, we measured HFA across all hippocampal electrodes while they examined hippocampal neurons with responses to old items that were significantly above baseline. Thus it is possible that these contrasting selection criteria contributed to our seemingly disparate findings.

Another important distinction is that we did not consider hippocampal subregions CA1 vs. CA3/DG separately, even though there is ample theoretical and empirical work suggesting these subregions may respond differently to these stimulus types, yet under certain circumstances both subregions can reflect activity consistent with separation or reinstatement (2, 16, 39, 41, 42, 44, 75–77). In the current study we were limited by the electrode placements, which were determined based on clinical criteria. Although our post-surgical images of electrode placements are not at a resolution to determine the hippocampal subregions of the electrodes, across participants approximately 60% of the electrodes were in the posterior portion of the hippocampus, which generally has a larger proportion of CA3/DG (78, 79). Future work remains to fully characterize how hippocampal subregions are modulated by experimental parameters such as task demands and stimulus types, as well as how these parameters impact the broader network of regions implicated in mnemonic discrimination.

It is critical to note that hippocampus was not the only brain region where HFA differentiated between similar and old trials. Like hippocampus, HFA in regions of the visual cortex (OTC) was also greater for similar compared to both old items and similar items classified as old. Interestingly, the HFA difference between similar items classified as old and correct similar items emerged in an earlier time window in OTC compared to hippocampus. The fact that activity in visual cortex differentiates highly similar stimuli is consistent with recent work showing that fMRI activity in inferior temporal regions can distinguish between highly similar stimuli (49, 80). Our results, using pattern analyses, also found evidence for mnemonic reinstatement during old item presentations. However, critically, unlike the hippocampus, OTC did not show any evidence for pattern separation. Rather, OTC patterns showed evidence for significant reinstatement, not separation, during similar item presentations. These results build on prior work implicating cortical reinstatement in memory success (33, 34, 81–83). It is important to note that the univariate and multivariate HFA results diverge in OTC: whereas univariate HFA was significantly greater for similar than old items, multivariate pattern analyses show that this may be related to reinstatement and not separation of the similar items.

It is tempting to think that pattern separation and reinstatement may be relatively automatic processes that support attention to overlapping and novel features of an environment during memory decision. Indeed, the fact that the pattern of neural effects were evident in OTC and DLPFC in both versions of the task (but not hippocampus, see above) suggests that the processes supporting memory discrimination in these regions are relatively automatic. OTC may be sensitive to the perceptual details in repeated items irrespective of the task. It is also intriguing to speculate that the early univariate effects in OTC may be related to a familiarity signal which has also extensively been shown to be rapid and relatively automatic (1, 84–86). Further work that specifically differentiates the subjective sense of familiarity from recollection, however, would be needed before this claim could be directly tested. In DLPFC, differences in univariate HFA paralleled those of OTC, with significantly greater HFA for similar than old items in both tasks. The univariate findings fit well with the reported role of DLPFC in both successful memory encoding (58–63) and retrieval (24, 43, 64, 65), as in the current study it is the case that similar items may require both more encoding of the novel details, while simultaneously placing greater demands on retrieval than old items. Our results are also consistent with the role of DLPFC in post-retrieval monitoring, a process by which the retrieved information is assessed based on task demands (64, 87–89).Our tasks arguably recruited such post-retrieval monitoring as similar and old items both required retrieval of the item’s original presentation, yet depending on the task and stimulus type would either lead to a response of ‘sim’ or ‘old’. Some studies have also reported that DLPFC activity differs based on task demands (65, 89–91), which may seem inconsistent with our findings of consistent DLPFC effects in both tasks.However, both of our tasks encouraged participants to retrieve a similar item’s matching first item, and thus the qualitative similarities in DLPFC activity across tasks may reflect how memory retrieval and post-retrieval monitoring were used in both task types.

In summary, our findings underscore the unique role of the hippocampus in mnemonic decisions. In the hippocampus, pattern analyses revealed significant pattern separation of items that were similar but not identical to an earlier item. By contrast, the hippocampus exhibited significant reinstatement of encoding activity during presentation of exact repeats. Reinstatement and separation in hippocampus occurred in the same late time window, suggesting that these processes may emerge over similar time scales. By contrast, an earlier visual region in OTC also exhibited reinstatement of old items, but exhibited reinstatement for similar items as well.Further, in this visual region, reinstatement occurred in an earlier time window than hippocampus. Taken together, these results provide support for the idea that the occipitotemporal and prefrontal cortical regions may be sensitive to the demands required to eaate highly similar lures from exact repeats – an especially challenging mnemonic operation – but that only the hippocampus may promote distinctive representations for these highly similar lures.

## Materials and Methods

### Participants

Five participants (4 female; 19-42 years old) with intractable epilepsy were recruited via the Comprehensive Epilepsy Center of the New York University School of Medicine. Participants had elected to undergo intracranial monitoring for clinical purposes, and provided informed consent to participate in this study under the approval of the local Institutional Review Board. Relevant clinical and demographic information for these participants is summarized in Table S1.

### Task design

Participants performed two separate blocks: one block of the fine-grain task and one block of the coarse-grain task (Fig. 1A,C). The order of the blocks was counterbalanced across participants. In each block, participants were presented with a series of images on a computer screen. Each image was either novel (‘new’), an exact repetition of a prior new stimulus (‘old’), or an image that was highly overlapping, but not identical to, a prior new stimulus (‘similar’). Each image was presented for 2.5-5s, with a blank 2.5s inter-stimulus interval separating trials.Presentation of the stimulus terminated following a participant’s response or 2.5s, whichever came later. If no response was made after 5s, item presentation ended.

In the fine-grain task block, participants were presented with 96 new images. Of these, half were presented again as old images and the other half were presented as similar trials. The number of intervening items between a new image and its subsequent old/similar trial varied between 1-8 trials. Participants were instructed to indicate, on each trial, whether the presented image was ‘new’, ‘old’, or ‘similar’.The three response options appeared in black on the bottom of the stimulus screen, in the same order as the response keys.

The coarse-grain task block had the same design as the fine-grain block, except that participants were instructed to designate the similar items as ‘old’. Thus, both actual repetitions and similar ‘repetitions’ could be designated as ‘old’, relieving the requirement to perform the fine-grained discrimination between actual old and similar items.

One participant only completed the task for 64 new items (and thus 32 similar and old items each) for each of the blocks.

### Electrophysiology

#### Recording

ECoG activity was recorded continuously during both blocks. Each participant had grid, strip and depth electrodes, and electrode placements were determined based on clinical criteria. ECoG activity was recorded with a custom built neural recording system (NSpike) at 10kHz. Sync pulses sent at the onset of each stimulus presentation and each participant response allowed for alignment of the ECoG data with trial onsets as well as behavioral responses.

#### Electrode localization

Hippocampal electrodes were manually identified with individual patient’s post-implantation magnetic resonance (MR) images using visual inspection of synchronized axial, coronal and sagittal slices (according to (92); see e.g. Fig. 1C,D). We first defined the posterior border by the first slice where gray matter appeared inferior and medial to the lateral ventricle. Then, moving anteriorly, wherever possible we used the landmarks of the lateral ventricle, white matter, and uncal recess to inform the borders. Across participants, electrodes were identified in the hippocampal head, body, and tail, yet there were insufficient electrodes to consider these subregions separately.

In addition to the hippocampus, we analyzed data from two additional regions: DLPFC and OTC. DLPFC electrodes were identified as any electrodes in the middle frontal gyrus (anterior to premotor cortex, i.e. Brodmann areas 9 and 46; (93)) based on visual inspection of each patient’s reconstructed 3D cortical surface using pre-and post-implantation MR scans (94). OTC electrodes were identified using a combination of the MR images and 3D brain reconstructions. This ROI was bounded superiorly by the inferior temporal sulcus and posteriorly by the occipital lobe, guided by the parieto-occipital fissure and the temporo-occipital incisures (93), anteriorly by the hippocampal tail (92), and ventrally by the occipitotemporal sulcus (95). The number of electrodes per region of interest is provided in Table S2.

#### Preprocessing

Data were downsampled to 300 Hz and each electrode was referenced to the mean activity across all of the patient’s electrodes, with the mean weighted such that each grid, strip or depth contributed equally (96). To remove electrical line noise, data were filtered at 60 Hz with a fourth order 2 Hz stopband Butterworth notch filter.

### Analysis

#### Conditions

When comparing a first presentation item (new) to a second presentation item (e.g. old), we consider the same set of stimuli in both cases: i.e. those items that were correctly classified as new for their first presentation and subsequently correctly classified as e.g. old items for the second presentation. In this way, the comparisons between first and second presentation items were matched for the number of observations and the types of stimuli that were tested. Furthermore, all comparisons between old and similar items were only conducted when their first presentations were correctly identified as ‘new’, thus providing some control for initial encoding.

#### High frequency activity (HFA)

Given that there were no clear peaks in the power spectrum in higher frequencies, we defined a high frequency activity (HFA) band at 45-115 Hz, above the beta band but below the second line noise harmonic. We calculated spectral power by applying a Morlet wavelet transform (wavelet number = 6) during stimulus presentation (0-2000ms post-stimulus onset) at 5 Hz intervals for each electrode and trial within an ROI. A 1000ms buffer was included on both sides of the data to minimize edge effects. Due to the broad distribution of power values we took the (natural) log transform of the power values. Power values for a particular presentation trial were normalized by subtracting the mean power at the same frequency during the corresponding baseline period 1500-500ms prior to stimulus onset.

#### HFA Univariate Power

Power was calculated as described above in the HFA band. Next, we calculated the mean power over time for four non-overlapping 500ms time bins. A Wilcoxon rank sum test was then used to compare pairs of conditions separately for each participant, electrode, frequency band and time bin. To determine significance between pairs of conditions, we used the summed Z method, an approach meant to assess significance with many observations per subjects but few subjects (62, 97, 98). With this approach, an empirical Z value is obtained from the experimental data (reported as the actual Z in the Results) and compared to a null distribution obtained using a permutation procedure (reported as the null mean Z in the Results, taken from 1000 random shuffles of the labels for each condition). The point at which the empirical Z score for a particular region fell in the null distribution determined the p value between conditions. Unless noted otherwise in the text, reported p values are Bonferroni corrected for the number of time windows. In addition, for illustrative purposes only in Figs 2, 5, 6, HFA in the top rows of is plotted as the mean across every 50ms with a 10ms sliding time window.

#### HFA spatiotemporal pattern similarity (STPS)

At each electrode for each trial, an HFA pattern vector was constructed for each 500ms time bin and type/response condition. Specifically, HFA was calculated at each electrode as above, in nonoverlapping 50ms time bins. In this way, 10 (50ms) time bins X the number of subject’s electrodes were included in each HFA pattern to yield a single vector of HFA values per trial.

To ensure that condition differences in pattern vectors do not reflect condition differences in univariate HFA, mean HFA across all trials of a stimulus/response type (e.g. correct old in the fine-grain task) was subtracted from each time-frequency element in every vector, within subject. These vectors could then be compared to each other to determine their correlation, or similarity.Matched pattern similarity was calculated as the Pearson’s r correlation between the spatiotemporal pattern vector an item’s first presentation (correct new) and the pattern vector of the item’s second presentation (as correct old, correct sim, incorrect sim). We used permutation tests to assess significance of pattern similarity values, as this allowed us to estimate a fair baseline of expected pattern similarity within and across participants. Specifically, we permuted the trial labels of the first presentations, and calculated the pattern similarity between first and second presentations based on these permuted trial labels at each time bin. The null distribution was defined as the mean of the pattern similarity values across 200 such permutations. The point at which the actual matched pattern similarity fell in the null distribution determined the p value, which was then Bonferroni corrected for multiple time windows. In a similar way, when comparing pairs of conditions, we calculated the difference between the values of the matched pairs and compared this to the difference between the values for the unmatched pairs in the null distribution, such that the point at which the actual matched difference fell on the null difference distribution determined the p values. In the Results, we report the mean STPS values from the empirical and null distributions.

### Correlation between univariate HFA and HFA STPS

For each participant, we then examined whether univariate HFA was related to STPS. Specifically, hippocampal HFA STPS was calculated for matched old-new pairs in each of the 500ms time bins. We then took the trial-by-trial correlation of the HFA pattern similarity difference values with HFA univariate activity during the same time bin as the STPS (i.e. for each of the 500ms bins). Next, for each participant we calculated an expected null distribution of correlation values by randomly shuffling the trial labels of HFA pattern similarity values, and taking the correlation of these shuffled pairs, for 200 shuffles of the data. The p value was determined by where the actual mean correlation fell on the mean shuffled distribution, and was Bonferroni corrected. In the Results, we report the mean STPS values from the empirical and null distributions.

### Data sharing

Data and analysis code will be made available upon completion of all planned analyses.

## Acknowledgments

We thank Thomas Thesen, Olga Felsovalyi and Amy Trongnetrpunya for their assistance with data collection, as well as the patients who participated in this study. This work was supported by MH106266 to Lynn Lohnas and MH074692 to Lila Davachi.

### Author contributions

K.D. and L.D. designed research; K.D., O.D. and W.K.D. performed research; L.J.L. analyzed data; and L.J.L., K.D. and L.D. wrote the paper.

## References

1. Eichenbaum H, Yonelinas AP, Ranganath C (2007) The medial temporal lobe and recognition memory. Annu Rev Neurosci 30:123–152.

2. Yassa MA, Stark CEL (2011) Pattern separation in the hippocampus. Trends Neurosci 34(10):515–525.

3. O’Reilly RC, Norman KA (2002) Hippocampal and neocortical contributions to memory: Advances in the complementary learning systems framework. Trends Cogn Sci 6(12):505–510.

4. Treves A, Rolls ET (1994) Computational analysis of the role of the hippocampus in memory. Hippocampus 4(3):374–391.

5. Hulbert JC, Norman KA (2015) Neural differentiation tracks improved recall of competing memories following interleaved study and retrieval practice. Cereb Cortex 25(10):3994–4008.

6. Favila SE, Chanales AJH, Kuhl BA (2016) Experience-dependent hippocampal pattern differentiation prevents interference during subsequent learning. Nat Commun 6:1–10.

7. Chanales AJH, Oza A, Favila SE, Kuhl BA (2017) Overlap among spatial memories triggers divergence of hippocampal representations. Curr Biol 27(15):1–47.

8. Marr D (1971) Simple memory: A theory for archicortex. Source Philos Trans R Soc London Ser B, Biol Sci 262(841):23–81.

9. Yassa MA, et al. (2011) Pattern separation deficits associated with increased hippocampal CA3 and dentate gyrus activity in nondemented older adults. Hippocampus 21(9):968–979.

10. Tompary A, Duncan K, Davachi L (2016) High-resolution investigation of memory-specific reinstatement in the hippocampus and perirhinal cortex. Hippocampus 26(8):995–1007.

11. Duncan K, Sadanand A, Davachi L (2012) Memory’s penumbra: episodic memory decisions induce lingering mnemonic biases. Science 337(6093):485–7.

12. Duncan K, Tompary A, Davachi L (2014) Associative Encoding and Retrieval Are Predicted by Functional Connectivity in Distinct Hippocampal Area CA1 Pathways. J Neurosci 34(34):11188–98.

13. Jezek K, Henriksen EJ, Treves A, Moser EI, Moser M-B (2011) Theta-paced flickering between place-cell maps in the hippocampus. Nature 478(7368):246–9.

14. Norman KA, O’Reilly RC (2003) Modeling hippocampal and neocortical contributions to recognition memory: a complementary-learning-systems approach. Psychol Rev 110(4):611–46.

15. Leutgeb S, et al. (2005) Independent codes for spatial and episodic memory in hippocampal neuronal ensembles. Science 309(5734):619–623.

16. Lee I, Yoganarasimha D, Rao G, Knierim JJ (2004) Comparison of population coherence of place cells in hippocampal subfields CA1 and CA3. Nature 430(6998):456–459.

17. Lee H, Wang C, Deshmukh SS, Knierim JJ (2015) Neural population evidence of functional heterogeneity along the CA3 transverse axis: Pattern completion versus pattern separation. Neuron 87(5):1093–1105.

18. McKenzie S, et al. (2014) Hippocampal representation of related and opposing memories develop within distinct, hierarchically organized neural schemas. Neuron 83(1):202–15.

19. Kriegeskorte N, Mur M, Bandettini PA (2008) Representational similarity analysis - connecting the branches of systems neuroscience. Front Syst Neurosci 2:1–28.

20. Backus AR, Bosch SE, Ekman M, Grabovetsky AV, Doeller CF (2016) Mnemonic convergence in the human hippocampus. Nat Commun 7(11991):1–9.

21. Mack ML, Preston AR (2016) Decisions about the past are guided by reinstatement of specific memories in the hippocampus and perirhinal cortex. Neuroimage 127:144–157.

22. van den Honert RN, McCarthy G, Johnson MK (2016) Reactivation during encoding supports the later discrimination of similar episodic memories. Hippocampus 26(9):1168–1178.

23. Staresina BP, et al. (2016) Hippocampal pattern completion is linked to gamma power increases and alpha power decreases during recollection. Elife 5:1–18.

24. Sederberg PB, et al. (2007) Gamma oscillations distinguish true from false memories. Psychol Sci 18(11):927–32.

25. Manning JR, Polyn SM, Baltuch GH, Litt B, Kahana MJ (2011) Oscillatory patterns in temporal lobe reveal context reinstatement during memory search. Proc Natl Acad Sci. doi:10.1073/pnas.1015174108.

26. Manning JR, Sperling MR, Sharan A, Rosenberg E a, Kahana MJ (2012) Spontaneously reactivated patterns in frontal and temporal lobe predict semantic clustering during memory search. J Neurosci 32(26):8871–8.

27. Yaffe RB, et al. (2014) Reinstatement of distributed cortical oscillations occurs with precise spatiotemporal dynamics during successful memory retrieval. Proc Natl Acad Sci U S A 111(52):18727–18732.

28. Yaffe RB, Shaikhouni A, Arai J, Inati SK, Zaghloul KA (2017) Cued memory retrieval exhibits reinstatement of high gamma power on a faster timescale in the left temporal lobe and prefrontal cortex. J Neurosci 37(17):3810–16.

29. Ritchey M, Wing EA, Labar KS, Cabeza R (2013) Neural similarity between encoding and retrieval is related to memory via hippocampal interactions. Cereb cortex:2818–2828.

30. Wing EA, Ritchey M, Cabeza R (2015) Reinstatement of individual past events revealed by the similarity of distributed activation patterns during encoding and retrieval. J Cogn Neurosci:1–10.

31. LaRocque KF, et al. (2013) Global similarity and pattern separation in the human medial temporal lobe predict subsequent memory. J Neurosci 33(13):5466–5474.

32. Staresina BP, Henson RN, Kriegeskorte N, Alink A (2012) Episodic reinstatement in the medial temporal lobe. J Neurosci 32(50):18150–18156.

33. Danker JF, Davachi L (2013) Cognitive neuroscience of episodic memory. The Oxford Handbook of Cognitive Neuroscience, eds Ochsner KN, Kosslyn S (Oxford University Press), pp 375–288.

34. Davachi L, Preston AR (2014) The medial temporal lobe and memory eds Gazzaniga MS, Mangun GR (MIT Press, Cambridge, MA). 5th Ed.

35. Danker JF, Tompary A, Davachi L (2016) Trial-by-trial hippocampal encoding activation predicts the fidelity of cortical reinstatement during subsequent retrieval. Cereb Cortex: 1–10.

36. Bosch SE, Jehee JFM, Fernández G, Doeller CF (2014) Reinstatement of Associative Memories in Early Visual Cortex Is Signaled by the Hippocampus. J Neurosci 34(22):7493–7500.

37. Gordon AM, Rissman J, Kiani R, Wagner AD (2014) Cortical reinstatement mediates the relationship between content-specific encoding activity and subsequent recollection decisions. Cereb Cortex 24(12):3350–3364.

38. St-Laurent M, Abdi H, Buchsbaum BR (2016) Distributed patterns of reactivation predict vividness of recollection. J Cogn Neurosci 27(10):2000– 2018.

39. Bakker A, Kirwan CB, Miller MB, Stark CEL (2008) Pattern separation in the human hippocampal CA3 and dentate gyrus. Science 319(5870):1640–2.

40. Kirwan CB, Stark CEL (2004) Medial temporal lobe activation during encoding and retrieval of novel face-name pairs. Hippocampus 14(7):919–30.

41. Lacy JW, Yassa MA, Stark SM, Muftuler LT, Stark CEL (2011) Distinct pattern separation related transfer functions in human CA3/dentate and CA1 revealed using high-resolution fMRI and variable mnemonic similarity. Learn Mem 18(1):15–18.

42. Duncan K, Ketz N, Inati SJ, Davachi L (2012) Evidence for area CA1 as a match/mismatch detector: a high-resolution fMRI study of the human hippocampus. Hippocampus 22(3):389–98.

43. Hannula DE, Ranganath C (2008) Medial temporal lobe activity predicts successful relational memory binding. J Neurosci 28(1):116–124.

44. Azab M, Stark SM, Stark CEL (2014) Contributions of human hippocampal subfields to spatial and temporal pattern separation. Hippocampus 24(3):293–302.

45. Copara MS, et al. (2014) Complementary roles of human hippocampal subregions during retrieval of spatiotemporal context. J Neurosci 34(20):6834–42.

46. Pidgeon LM, Morcom AM (2016) Cortical pattern separation and item-specific memory encoding. Neuropsychologia 85.

47. Leutgeb S, Leutgeb JK, Treves A, Moser M-B, Moser EI (2004) Distinct ensemble codes in hippocampal areas CA3 and CA1. Science 305(5688):1295– 1298.

48. Hashimoto R, et al. (2012) Changing the criteria for old/new recognition judgments can modulate activity in the anterior hippocampus. Hippocampus 22(2):141–148.

49. Kirwan CB, Stark CEL (2007) Overcoming interference: an fMRI investigation of pattern separation in the medial temporal lobe. Learn Mem 14(9):625–633.

50. Duncan K, Sadanand A, Davachi L (2012) Memory’s penumbra : Episodic memory decisions induce lingering mnemonic biases. Science 337(6093):485–7.

51. Fries P, Reynolds JH, Rorie AE, Desimone R (2001) Modulation of oscillatory neuronal synchronization by selective visual attention. Science 291(5508):1560–1563.

52. Hirabayashi T, et al. (2014) Distinct neuronal interactions in anterior inferotemporal areas of macaque monkeys during retrieval of object association memory. J Neurosci 34(28):9377–9388.

53. Manning JR, Jacobs J, Fried I, Kahana MJ (2009) Broadband ahifts in local field potential power spectra are correlated with single-neuron spiking in humans. J Neurosci 29(43):13613–13620.

54. Crone NE, Korzeniewska A, Franaszczuk PJ (2011) Cortical gamma responses: Searching high and low. Int J Psychophysiol 79(1):9–15.

55. Lachaux J-P, Axmacher N, Mormann F, Halgren E, Crone NE (2012) High-frequency neural activity and human cognition: past, present and possible future of intracranial EEG research. Prog Neurobiol 98(3):279–301.

56. Janowsky JS, Shimamura AP, Squire LR (1989) Source memory impairment in patients with frontal lobe lesions. Neuropsychologia 27(8):1043–1056.

57. McAndrews MP, Milner B (1991) The frontal cortex and memory for temporal order. Neuropsychologia 29(9):849–859.

58. Ranganath C, Johnson MK, D’Esposito M (2000) Left anterior prefrontal activation increases with demands to recall specific perceptual information. J Neurosci 20:1–5.

59. Jenkins LJ, Ranganath C (2010) Prefrontal and medial temporal lobe activity at encoding predicts temporal context memory. J Neurosci 30(46):15558–65.

60. Blumenfeld RS, Parks CM, Yonelinas AP, Ranganath C (2011) Putting the pieces together: the role of dorsolateral prefrontal cortex in relational memory encoding. J Cogn Neurosci 23(1):257–65.

61. Ranganath C, Cohen MX, Brozinsky CJ (2005) Working memory maintenance contributes to long-term memory formation: neural and behavioral evidence. J Cogn Neurosci 17(7):994–1010.

62. Sederberg PB, et al. (2007) Hippocampal and neocortical gamma oscillations predict memory formation in humans. Cereb Cortex (1190–1196).

63. Ranganath C, Johnson MK, D’Esposito M (2003) Prefrontal activity associated with working memory and episodic long-term memory. Neuropsychologia 41(3):378–389.

64. Dobbins IG, Foley H, Schacter DL, Wagner AD (2002) Executive control during episodic retrieval: multiple prefrontal processes subserve source memory. Neuron 35:989–996.

65. Ranganath C, Cohen MX, Dam C, D’Esposito M (2004) Inferior temporal, prefrontal, and hippocampal contributions to visual working memory maintenance and associative memory retrieval. J Neurosci 24(16):3917–3925.

66. Grill-Spector K, Henson RN, Martin A (2006) Repetition and the brain: neural models of stimulus-specific effects. Trends Cogn Sci 10(1):14–23.

67. Rodriguez Merzagora A, et al. (2014) Repeated stimuli elicit diminished high-gamma electrocorticographic responses. Neuroimage 85 Pt 2:844–52.

68. Zhang H, et al. (2015) Gamma power reductions accompany stimulus-specific representations of dynamic events. Curr Biol 25(5):635–640.

69. Gruber T, Müller MM (2005) Oscillatory brain activity dissociates between associative stimulus content in a repetition priming task in the human EEG. Cereb Cortex 15(1):109–116.

70. Rugg MD, et al. (2012) Item memory, context memory and the hippocampus: fMRI evidence. Neuropsychologia 50(13):3070–9.

71. Nolde SF, Johnson MK, Raye CL (1998) The role of prefrontal cortex during tests of episodic memory. Trends Cogn Sci 2(10):399–406.

72. Rugg MD, Yonelinas AP (2003) Human recognition memory: A cognitive neuroscience perspective. Trends Cogn Sci 7(7):313–319.

73. Merkow MB, Burke JF, Kahana MJ (2015) The human hippocampus contributes to both the recollection and familiarity components of recognition memory. Proc Natl Acad Sci 112(46):14378–14383.

74. Suthana NA, et al. (2015) Specific responses of human hippocampal neurons are associated with better memory. Proc Natl Acad Sci: 201423036.

75. Schapiro AC, Kustner L V, Turk-Browne NB (2012) Shaping of object representations in the human medial temporal lobe based on temporal regularities. Curr Biol 22(17):1622–7.

76. Wills TJ, Lever C, Cacucci F, Burgess N, O’Keefe J (2005) Attractor dynamics in the hippocampal representation of the local environment. Science 308(5723):873–876.

77. Leutgeb JK, Leutgeb S, Moser M-B, Moser EI (2007) Pattern separation in the dentate gyrus and CA3 of the hippocampus. Science 315(5814):961–966.

78. Malykhin N V., Lebel RM, Coupland NJ, Wilman AH, Carter R (2010) In vivo quantification of hippocampal subfields using 4.7 T fast spin echo imaging. Neuroimage 49(2):1224–1230.

79. Poppenk J, Evensmoen HR, Moscovitch M, Nadel L (2013) Long-axis specialization of the human hippocampus. Trends Cogn Sci:1–11.

80. Reagh ZM, Yassa M a (2014) Object and spatial mnemonic interference differentially engage lateral and medial entorhinal cortex in humans. Proc Natl Acad Sci U S A. doi:10.1073/pnas.1411250111.

81. Wheeler ME, Petersen SE, Buckner RL (2000) Memory’s echo: Vivid remembering reactivates sensory-specific cortex. Proc Natl Acad Sci 97(20):11125–11129.

82. Nyberg L, Habib R, McIntosh AR, Tulving E (2000) Reactivation of encoding-related brain activity during memory retrieval. Proc Natl Acad Sci 97:11120– 11124.

83. Polyn SM, Natu VS, Cohen JD, Norman KA (2005) Category-specific cortical activity precedes retrieval during memory search. Science 310(5756):1963–6.

84. Yonelinas AP (2002) The nature of recollection and familiarity: A review of 30 years of research. J Mem Lang 46(3):441–517.

85. Brown MW, Aggleton JP (2001) Recognition memory: What are the roles of the perirhinal cortex and hippocampus? Nat Rev Neurosci 2(1):51–61.

86. Yonelinas AP, Jacoby LL (2012) The process-dissociation approach two decades later: Convergence, boundary conditions, and new directions. Mem Cognit 40(5):663–680.

87. D’Esposito M, et al. (1995) The neural basis of the central executive system of working memory. Nature.

88. Henson RN, Shallice T, Dolan RJ (1999) Right prefrontal cortex and episodic memory retrieval: A functional MRI test of the monitoring hypothesis. Brain 122:1367–1381.

89. MacDonald AW, Cohen JD, Stenger VA, Carter CS (2000) Dissociating the role of the dorsolateral prefrontal and anterior cingulate cortex in cognitive control. Science 288(June):1835–1838.

90. Braver TS, et al. (1997) A parametric study of prefrontal cortex involvement in human working memory. Neuroimage 5(1):49–62.

91. Rudorf S, Hare TA (2014) Interactions between dorsolateral and ventromedial prefrontal cortex underlie context-dependent stimulus valuation in goal-directed choice. J Neurosci 34(48):15988–15996.

92. Pruessner JC, et al. (2000) Volumetry of hippocampus and amygdala with high-resolution MRI and three-dimensional analysis software: Minimizing the discrepancies between laboratories. Cereb Cortex 10(4):433–442.

93. Duvernoy HM (1991) The human brain: Surface, blood supply, and three-dimensional sectional anatomy (Springer). 2nd Ed.

94. Yang AI, et al. (2012) Localization of dense intracranial electrode arrays using magnetic resonance imaging. Neuroimage 63(1):157–65.

95. Grill-Spector K, Weiner KS (2013) The functional architecture of the ventral temporal cortex and its role in categorization. 18(9):1199–1216.

96. Serruya MD, Sederberg PB, Kahana MJ (2014) Power shifts track serial position and modulate encoding in human episodic memory. Cereb Cortex 24(2):403–13.

97. Sederberg PB, et al. (2006) Oscillatory correlates of the primacy effect in episodic memory. Neuroimage 32(3):1422–31.

98. Sederberg PB, Kahana MJ, Howard MW, Donner EJ, Madsen JR (2003) Theta and gamma oscillations during encoding predict subsequent recall. J Neurosci 23(34):10809–14.

